# Reconciling high-throughput gene essentiality data with metabolic network reconstructions

**DOI:** 10.1101/415448

**Authors:** Anna S. Blazier, Jason A. Papin

## Abstract

The identification of genes essential for bacterial growth and survival represents a promising strategy for the discovery of antimicrobial targets. Essential genes can be identified on a genome-scale using transposon mutagenesis approaches; however, variability between screens and challenges with interpretation of essentiality data hinder the identification of both condition-independent and condition-dependent essential genes. To illustrate the scope of these challenges, we perform a large-scale comparison of multiple published *Pseudomonas aeruginosa* gene essentiality datasets, revealing substantial differences between the screens. We then contextualize essentiality using genome-scale metabolic network reconstructions and demonstrate the utility of this approach in providing functional explanations for essentiality and reconciling differences between screens. Genome-scale metabolic network reconstructions also enable a high-throughput, quantitative analysis to assess the impact of media conditions on the identification of condition-independent essential genes. Our computational model-driven analysis provides mechanistic insight into essentiality and contributes novel insights for design of future gene essentiality screens and the identification of core metabolic processes.

**Author Summary:** With the rise of antibiotic resistance, there is a growing need to discover new therapeutic targets to treat bacterial infections. One attractive strategy is to target genes that are essential for growth and survival. Essential genes can be identified with transposon mutagenesis approaches; however, variability between screens and challenges with interpretation of essentiality data hinder the identification and analysis of essential genes. We performed a large-scale comparison of multiple gene essentiality screens of the microbial pathogen *Pseudomonas aeruginosa*. We implemented a computational model-driven approach to provide functional explanations for essentiality and reconcile differences between screens. The integration of computational modeling with high-throughput experimental screens may enable the identification of drug targets with high-confidence and provide greater understanding for the development of novel therapeutic strategies.

## Introduction

With the rise of antibiotic resistance, there is a growing need to discover new therapeutic targets to treat bacterial infections. One attractive strategy is to target genes that are essential for growth and survival [1–4]. Discovery of such genes has been a long-standing interest, and advances in transposon mutagenesis combined with high-throughput sequencing have enabled their identification on a genome-scale. Transposon mutagenesis screens have been used to discriminate between *in vivo* and *in vitro* essential genes [1,5], discover genes uniquely required at different infection sites [6], and assess the impact of co-infection on gene essentiality status [7]. However, nuanced differences in experimental methods and data analysis can lead to variable essentiality calls between screens and hamper the identification of essential genes with high-confidence [8,9]. Additionally, a central challenge of these screens is in interpreting why a gene is or is not essential in a given condition, hindering the identification of promising drug targets.

These data are often used to validate and curate genome-scale metabolic network reconstructions (GENREs) [10,11]. GENREs are knowledgebases that capture the genotype-to-phenotype relationship by accounting for all the known metabolic genes and associated reactions within an organism of interest. These reconstructions can be converted into mathematical models and subsequently used to probe the metabolic capabilities of an organism or cell type in a wide range of conditions. GENREs of human pathogens have been used to discover novel drug targets [12], determine metabolic constraints on the development of antibiotic resistance [13], and identify metabolic determinants of virulence [14]. Importantly, GENREs can be used to assess gene essentiality by simulating gene knockouts. Through *in silico* gene essentiality analysis, GENREs can be useful in the systematic comparison of gene essentiality datasets.

Here, we perform the first large-scale, comprehensive comparison and reconciliation of multiple gene essentiality screens and contextualize these datasets using genome-scale metabolic network reconstructions. We apply this framework to the Gram-negative, multi-drug resistant pathogen *Pseudomonas aeruginosa*, using several published transposon mutagenesis screens performed in various media conditions and the recently published GENREs for strains PAO1 and PA14. We demonstrate the utility of interpreting transposon mutagenesis screens with GENREs by providing functional explanations for essentiality, resolving differences between the screens, and highlighting gaps in our knowledge of *P. aeruginosa* metabolism. Finally, we perform a high-throughput, quantitative analysis to assess the impact of media conditions on identification of core essential genes. This work demonstrates how genome-scale metabolic network reconstructions can help interpret gene essentiality data and guide future experiments to further enable the identification of essential genes with high-confidence.

## Results

### Comparison of candidate essential genes reveals variability across transposon mutagenesis screens

We obtained data from several published transposon mutagenesis screens for *P. aeruginosa* strains PAO1 and PA14 in various media conditions and determined candidate essential genes for each screen as described in Methods (Table S1) [15–19]. Briefly, where available, we used the published essential gene lists identified by the authors of the screen. Otherwise, we defined genes as essential in a particular screen if the corresponding mutant did not appear in that screen, suggesting that a mutation in the corresponding gene resulted in a non-viable mutant. Candidate essential gene lists ranged in size from 179 to 913 for PAO1 and from 510 to 1544 for PA14, suggesting substantial variability between the screens (Table 1, Dataset_S1, Dataset_S2). To investigate the similarity between the different candidate essential gene lists for the two strains, we performed hierarchical clustering with complete linkage on the dissimilarity between the candidate essential gene lists, as measured by Jaccard distance (Figure 1A and 1C). Interestingly, the screens clustered by publication rather than by media condition for both strains. As an example from the PAO1 screens, rather than clustering by lysogeny broth (LB) media, sputum media, pyruvate minimal media, and succinate minimal media, all three of the screens from the Lee et al. publication clustered together, all three of the screens analyzed in the Turner et al. publication clustered together, and the Jacobs et al. transposon mutant library clustered independently. This result suggests that experimental technique and downstream data analysis play a large role in determining essential gene calls, motivating the importance of comparing several screens to identify consensus essential gene lists, or genes identified as essential across multiple screens.

**Table 1.**
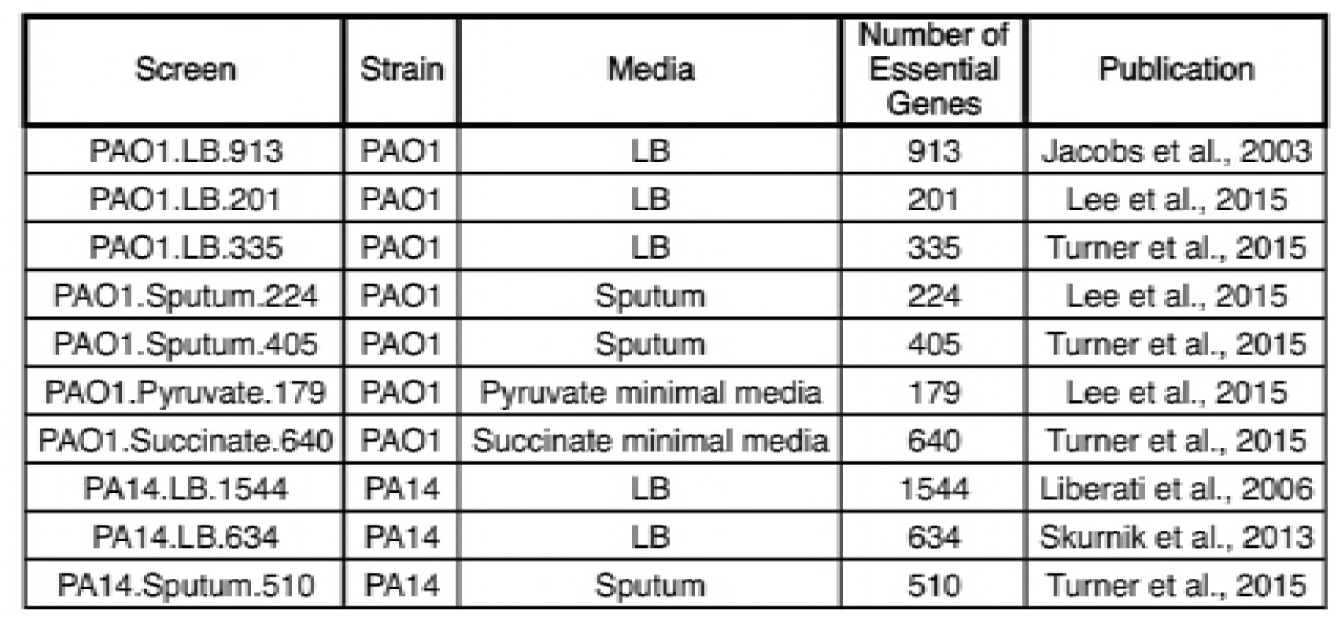
Characteristics of the *in vitro* transposon mutagenesis screens.

| Screen | Strain | Media | Number of Essential Genes | Publication |
| --- | --- | --- | --- | --- |
| PAO1.LB.913 | PAO1 | LB | 913 | Jacobs et al., 2003 |
| PAO1.LB.201 | PAO1 | LB | 201 | Lee et al., 2015 |
| PAO1.LB.335 | PAO1 | LB | 335 | Turner et al., 2015 |
| PAO1.Sputum.224 | PAO1 | Sputum | 224 | Lee et al., 2015 |
| PAO1.Sputum.405 | PAO1 | Sputum | 405 | Turner et al., 2015 |
| PAO1.Pyruvate.179 | PAO1 | Pyruvate minimal media | 179 | Lee et al., 2015 |
| PAO1.Succinate.640 | PAO1 | Succinate minimal media | 640 | Turner et al., 2015 |
| PA14.LB.1544 | PA14 | LB | 1544 | Liberati et al., 2006 |
| PA14.LB.634 | PA14 | LB | 634 | Skurnik et al., 2013 |
| PA14.Sputum.510 | PA14 | Sputum | 510 | Turner et al., 2015 |

**Figure 1.**
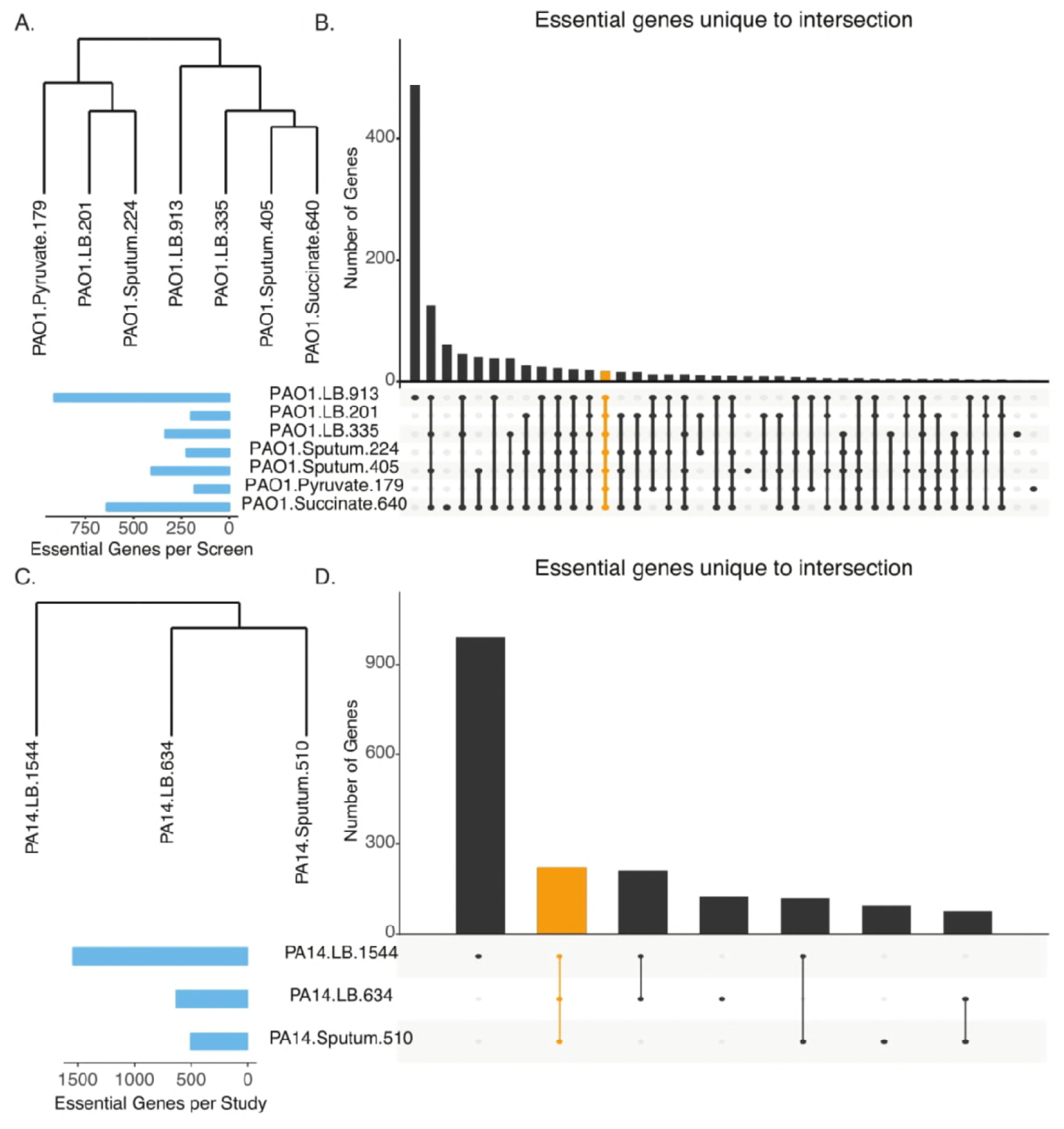
Comparison of candidate essential genes from transposon mutagenesis screens reveals variability. (A and C). Hierarchical clustering of candidate essential gene lists from transposon mutagenesis screens for PAO1 and PA14, respectively. (B and D). Overlap analysis of candidate essential gene lists for transposon mutagenesis screens for PAO1 and PA14, respectively. Blue bars indicate the total number of candidate essential genes identified in each screen. Black bars indicate the number of candidate essential genes unique to the intersection given by the filled-in dots. The orange bar indicates the overlap for all screens for either PAO1 (Panel B) or PA14 (Panel D). For the relationship between the overlap analysis and venn diagrams, see Figures S1 and S2.

We then measured the overlap of the candidate essential gene lists to calculate how many genes were shared across all the screens as well as those unique to particular sets of screens, defined as intersections (Figure 1B and 1D). For both strains, the candidate essential genes unique to the transposon mutant libraries (i.e., PAO1.LB.913 and PA14.LB.1544) accounted for the largest grouping, reflecting the disproportionately large size of both screens’ candidate essential gene lists relative to the transposon sequencing screens. Approximately 63% and 54% of the essential genes were unique to the PA14.LB.1544 and PAO1.LB.913 screens, respectively. While genes were uniquely essential for PAO1 on individual LB screens, there were no genes uniquely essential to all three LB screens; rather, the genes identified as commonly essential in all three LB screens were also identified in one or more of the sputum, pyruvate and succinate screens. This trend also held for the PAO1 sputum screens; however, 61 genes were uniquely identified in the succinate minimal media screen and two genes were uniquely identified in the pyruvate minimal media screen, perhaps reflecting the more stringent conditions of the minimal media screens relative to the more rich conditions of the LB and sputum screens.

This analysis revealed substantial differences in the overlap of the candidate essential genes across the screens. Using the number of intersections as an indicator of variability, comparison of the PAO1 screens resulted in more than 30 intersections, while comparison of the PA14 screens resulted in seven, highlighting the discrepancies between the screens for both *P. aeruginosa* strains. This heterogeneity across the screens could be attributed to a number of factors such as screening approach (e.g., individually mapped mutants versus transposon sequencing), library complexity, metrics of essentiality, data analysis, and the media conditions tested. To investigate the possibility that these discrepancies were completely due to data analysis alone and not experimental differences, we re-analyzed the sequencing data for the PAO1 transposon sequencing screens performed on LB where sequencing data was publicly available using the same analytical pipeline (Figure S1)[18,20]. As expected, when the same analysis pipeline was applied to the two screens, there was an increase in the number of commonly essential genes compared to the overlap between the published results. However, there were still genes that were identified as uniquely essential to each screen. These results suggest that differences in data processing alone do not account for the observed variability between the screens but that experimental differences, such as library complexity, number of replicates, and read depth, likely also contribute.

To determine potential core essential genes (i.e., genes that are essential regardless of media or other conditions), we measured the number of genes that were shared by all of the screens for either PAO1 or PA14. Surprisingly, only 17 genes were shared by all PAO1 screens while 192 genes were shared by all PA14 screens. These numbers of core essential genes are lower than expected, particularly for strain PAO1. Typically, essential genes are thought to number a few hundred for the average bacterial genome [21]. We reasoned that this unexpectedly low number of observed core essential genes might be due to the variety of media conditions across the PAO1 screens, so we repeated our analysis focusing only on the LB media screens for both PA14 and PAO1 (Figure S2). Interestingly, the trends remained the same, with 434 genes shared across both PA14 LB media screens and only 44 genes shared across all PAO1 LB media screens. Overall, the PA14 screens had higher numbers of essential genes compared to those for PAO1, with all the PA14 screens having at least 400 essential genes. In contrast, there were four PAO1 screens with less than 350 essential genes. Together, these differences suggest greater variability for transposon mutagenesis in PAO1 compared to PA14. Strain-specific differences in essentiality have been reported previously but are underappreciated [22]. This result adds to the growing literature emphasizing how the genetic background of the strain analyzed may impact the identification of essential genes. Nevertheless, the identified core essential genes point to genes that may potentially be indispensable for bacterial growth and survival regardless of condition.

Taken together, results from this comparison revealed vast differences between the candidate essential gene lists across screens, even for those from the same media condition. These differences may be due to a number of factors such as experimental screening approach, library complexity, read depth, and downstream data analysis. Ultimately, this variability complicates the discovery of essential genes with high-confidence.

### Contextualization of gene essentiality datasets using genome-scale metabolic network reconstructions

A central challenge of transposon mutagenesis screens lies in the interpretation of why a gene is or is not essential in a given condition. Here, we demonstrate the utility of genome-scale metabolic network reconstructions to contextualize gene essentiality and provide mechanistic explanations for the essentiality status of metabolic genes. To do this, we compared the *in vitro* candidate essential gene lists to predicted essential genes from the PAO1 and PA14 GENREs [23]. These GENREs were previously shown to predict gene essentiality with an accuracy of 91% [23]. For both models, we simulated *in silico* gene knockouts under media conditions that approximated those used in the *in vitro* screens and assessed the resulting impact on biomass synthesis as an approximation for growth (Dataset_S3, Dataset_S4). Genes were predicted to be essential if biomass production for the associated mutant model was below a standard threshold. Predicted essential gene lists for both the PAO1 and PA14 models under the different media conditions were compared to the candidate essential gene lists for each of the experimental screens and the matching accuracy between model predictions and the *in vitro* screens was assessed (Figure 2A, Table S2).

**Figure 2.**
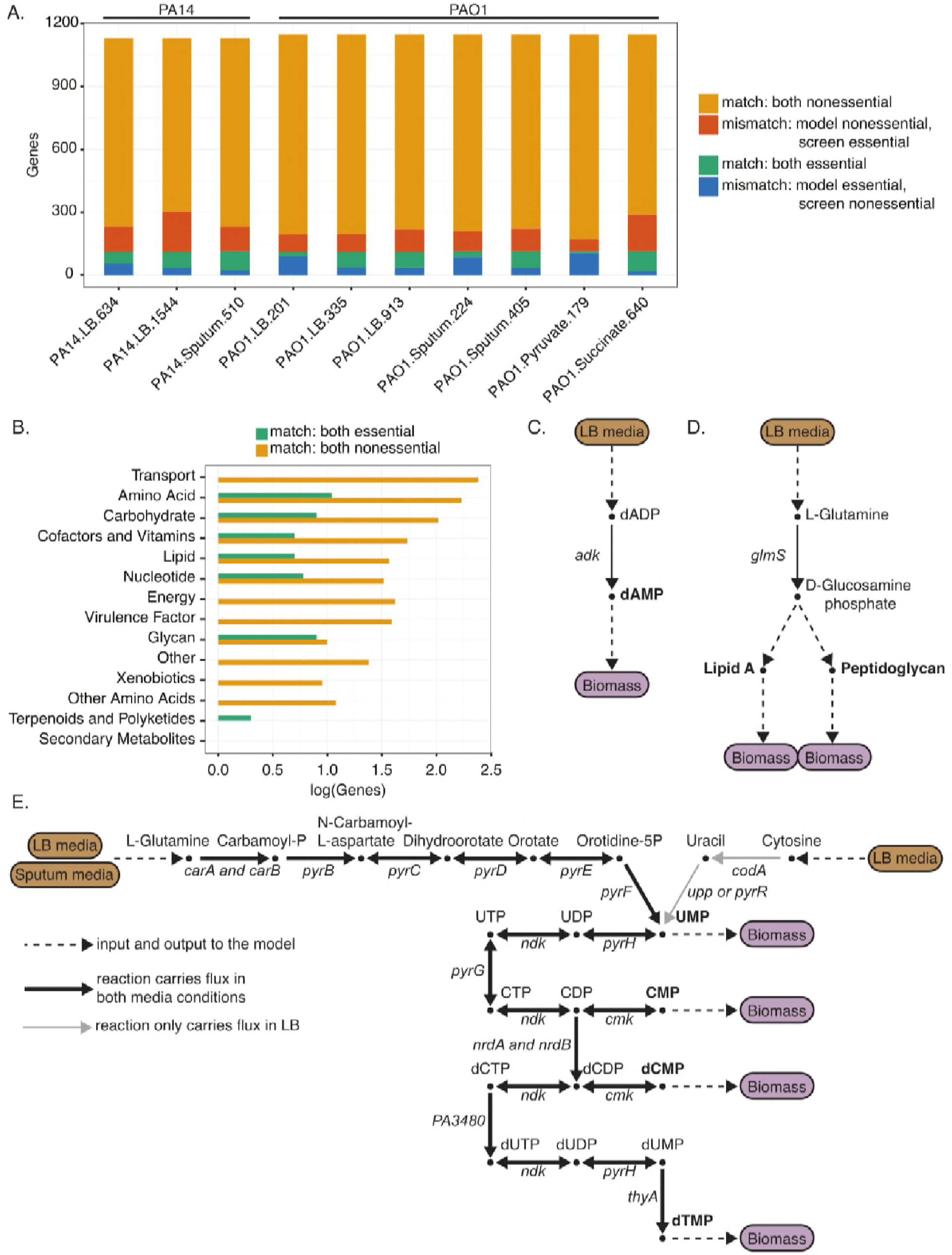
Contextualization of gene essentiality datasets using genome-scale metabolic network reconstructions. (A) Comparison of model essentiality predictions to *in vitro* essentiality screens. *In silico* gene knockouts were performed for both PA14 and PAO1 genome-scale metabolic network reconstructions to predict essential genes. Model-predicted essential genes were compared to the candidate essential genes for each *in vitro* screen. The bars show the result of this comparison, with orange indicating the number of genes for which both the model and experimental screen identified the gene as nonessential (match: both nonessential), red indicating the number of genes for which the model identified the gene as nonessential whereas the screen identified the gene as essential (mismatch: model-nonessential, screen-essential), green indicating the number of genes for which both the model and experimental screen identified the gene as essential (match: essential), and blue indicating the number of genes for which the model identified the gene as essential whereas the screen identified the gene as nonessential (mismatch: model-essential, screen-nonessential). (B) Functional subsystems for PA14 consensus essential and nonessential genes that were also correctly predicted to be essential or nonessential in the PA14 GENRE. Consensus essential and nonessential genes were identified for PA14 by comparing all three LB screens and determining genes essential or nonessential in all three screens. (C and D). Metabolic pathways demonstrating essentiality for the consensus essential genes *adk* and *glmS*, respectively. Dashed lines represent inputs and outputs of the pathway, or, as in D, multiple steps. Brown boxes indicate media inputs, while purple boxes indicate biomass outputs. Metabolites are labeled beside the nodes, with bold metabolites indicating biomass components. Genes associated with the specific reaction are indicated. (E). Flux activity in pyrimidine metabolism under both sputum and LB media conditions. Consensus LB essential genes were compared to consensus sputum essential genes for PAO1. The PAO1 GENRE was used to explain differences in essentiality between the two media-types. Black lines indicate that the reaction is capable of carrying flux under both sputum and LB conditions, while the gray lines indicate that the reaction does not carry flux in sputum conditions but does in LB conditions. Brown boxes are media inputs, purple boxes are biomass outputs. Metabolites are labeled above the nodes, with bold metabolites indicating biomass components. Many of these metabolites are involved in many reactions beyond pyrimidine metabolism. Gene-protein-reaction relationships are indicated in italics beside each reaction edge.

As expected, most genes were identified as nonessential by both the screens and the models. These nonessential genes likely encode redundant features in the metabolic network, such as isozymes or alternative pathways, or are involved in accessory metabolism, such as the production of small molecule virulence factors. Interestingly, the number of screen-essential genes predicted as nonessential was significantly larger than the number of screen-nonessential genes predicted as essential (p < 0.01, as measured by Wilcoxon signed-rank test). We hypothesize that the reason for this difference is due to the increased likelihood of an *in vitro* screen missing a gene, potentially due to gene length or transposition cold spots [16], and subsequently incorrectly identifying it as essential.

This analysis can help to provide specific functional explanations for essentiality. Where there is agreement between the model predictions and *in vitro* screens, we can use the network to explain why a gene is or is not essential. Similarly, we can analyze the network to explain why a gene may be essential in one media condition versus another. A mismatch denotes some discrepancy between the model predictions and the experimental results. These mismatches may point to a gap in the model, indicating that it is missing some relevant biological information. Alternatively, the mismatches may be due to experimental variability such as differences in environmental conditions or technique.

To begin contextualizing the gene essentiality datasets using the GENREs, we focused on metabolic genes that were identified as essential or as nonessential in all LB screens for either PAO1 or PA14 (which we termed “consensus essential genes” and “consensus nonessential genes”, respectively) (Table S3, Dataset_S5, Dataset_S6). Consensus essential genes have a greater likelihood of being truly essential rather than experimental artifacts since they were identified as such in multiple independent screens. We then compared these lists of consensus essential genes and consensus nonessential genes to the model predictions of essentiality in LB media.

From this comparison, we found 45 of 113 consensus essential genes predicted to be essential by the PA14 model and 777 of 800 consensus nonessential genes predicted to be nonessential by the PA14 model. For PAO1, we found seven of 15 consensus essential genes predicted to be essential by the PAO1 model and 843 of 863 consensus nonessential genes predicted as nonessential by the PAO1 model (Table S3). The low number of consensus essential genes for PAO1 reflects the high variability between screens, as highlighted in Figures 1 and S1.

We then used the models to delineate subsystem assignments for the model-predicted consensus essential and nonessential genes (Figure 2B for PA14 and Figure S3 for PAO1). As expected, the consensus nonessential genes spanned most subsystems within the network, likely due to redundancy in the network as well as the presence of accessory metabolic functions that are not critical for biomass production. In contrast, for PA14, the consensus essential genes were limited to seven of the 14 subsystems within the network (note that this trend does not hold for PAO1 because there were very few consensus essential genes to consider). These seven subsystems capture metabolic pathways that are critical for bacterial growth and survival. For instance, lipid metabolism is essential for building and maintaining cell membranes, while carbohydrate metabolism is critical for ATP generation. None of the genes involved in transport were consensus essential genes. Because we only considered screens performed in LB media, transport of individual important metabolites, such as a specific carbon sources, was not a limiting factor given the abundant availability of such compounds in rich media conditions. However, we would expect that if we considered screens performed under minimal media conditions, relevant transport genes would be essential for bacterial growth.

Because these consensus essential genes were also predicted to be essential by the model, we can use the network to provide functional reasons for essentiality. For example, both the model and screens identified the gene *adk*, encoding adenylate kinase, as essential. Using the model, we determined that when *adk* is not functional, the conversion of deoxyadenosine diphosphate (dADP) to deoxyadenosine monophosphate (dAMP) cannot proceed, impacting the cell’s ability to synthesize DNA and ultimately produce biomass (Figure 2C). The model can also tease out less obvious relationships. For instance, both the model and the screens identified *glmS*, encoding glucosamine-fructose-6-phosphate aminotransferase, as essential. Using the model, we found that when *glmS* is not functional, the conversion of L-Glutamine to D-Glucosamine phosphate cannot proceed. D-Glucosamine phosphate is an essential precursor to both Lipid A, a component of the endotoxin lipopolysaccharide, and peptidoglycan, which forms the cell wall (Figure 2D). For each of the model-predicted consensus essential genes, we identified which biomass components could not be synthesized when the gene was removed from the model (Dataset_S7 and Dataset_S8). Further analysis is necessary to tease out the metabolic pathways that prevent synthesis of these biomass metabolites; however, from the examples above it is evident that GENREs can provide both obvious and non-obvious functional explanations for essentiality, streamlining the interpretation of transposon mutagenesis screens.

In addition to identifying consensus essential and nonessential genes that were in agreement with the models, we also uncovered discrepancies between model predictions and experimental results. For PAO1 and PA14, respectively, there were 8 and 68 consensus essential genes that the models predicted to be nonessential and 20 and 23 consensus nonessential genes that the models predicted to be essential. These mismatches between model predictions and experimental results provide insight into gaps in our understanding of *P. aeruginosa* metabolism.

In the case where a consensus essential gene was predicted to be non-essential by the model, this result indicates that the model has some additional functionality that is not available *in vitro*. This result could be an inaccuracy of the network reconstruction or it could be a result of using a non-condition-specific network where the model has access to all possible reactions in the network. Because cells undergo varying states of regulation, gene essentiality can be modulated as a result. Thus, profiling data such as transcriptomics could be integrated into the network reconstruction to generate a condition-specific model to improve model predictions under specified conditions [24,25].

In contrast, in the case where a consensus nonessential gene was predicted to be essential, this result indicates that the model is missing key functionality, pointing to areas of potential model curation. Using this list of discrepancies to guide curation (Table 2), we performed an extensive literature review and found several suggested changes to the metabolic network reconstruction (Dataset_S9). For instance, we incorrectly predicted as essential the gene *fabI* (PA1806), which is linked to triclosan resistance; however, a recent study discovered an isozyme of *fabI* in PAO1 called *fabV* (PA2950) [26]. To account for this new information, we suggest changing the gene-protein-reaction (GPR) relationship for the 28 reactions governed by *fabI* to be “*fabI* OR *fabV*”, making *fabI* no longer essential in the model. Additionally, our model incorrectly predicted the genes *ygiH* (PA0581) and *plsX* (PA2969) to be essential due to a GPR formulation of “*ygiH* AND *plsX*” for several reactions in glycerolipid metabolism. Literature evidence suggests that the gene-product of *plsB* (PA3673) is also able to catalyze these reactions. Specifically, the gene-products of both *plsB* and the *ygiH*/*plsX* system are able to carry out the acylation of glycerol-3-phosphate from an acyl carrier protein whereas only the gene-product of *plsB* is able to carry out this reaction for acyl-CoA thioesters [27,28]. This experimental evidence motivates changing the GPRs for 16 reactions in glycerolipid metabolism.

**Table 2.**
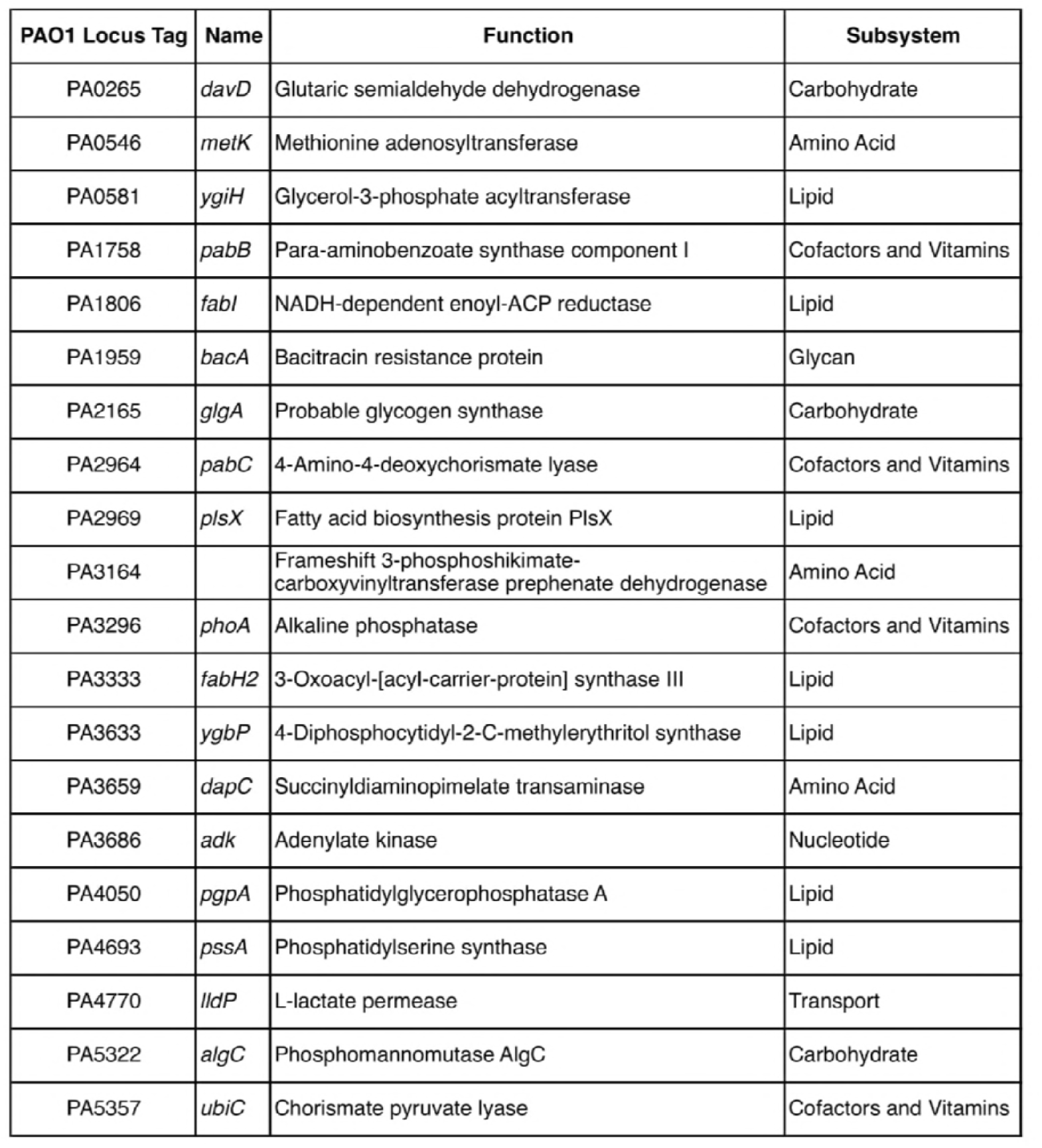
Discrepancies between model predicted essential genes and *in vitro* identified consensus nonessential genes for PAO1.

| PAO1 Locus Tag | Name | Function | Subsystem |
| --- | --- | --- | --- |
| PA0265 | *davD* | Glutaric semialdehyde dehydrogenase | Carbohydrate |
| PA0546 | *metK* | Methionine adenosyltransferase | Amino Acid |
| PA0581 | *ygiH* | Glycerol-3-phosphate acyltransferase | Lipid |
| PA1758 | *pabB* | Para-aminobenzoate synthase component I | Cofactors and Vitamins |
| PA1806 | *fabI* | NADH-dependent enoyl-ACP reductase | Lipid |
| PA1959 | *bacA* | Bacitracin resistance protein | Glycan |
| PA2165 | *glgA* | Probable glycogen synthase | Carbohydrate |
| PA2964 | *pabC* | 4-Amino-4-deoxychorismate lyase | Cofactors and Vitamins |
| PA2969 | *plsX* | Fatty acid biosynthesis protein PlsX | Lipid |
| PA3164 |  | Frameshift 3-phosphoshikimate-carboxyvinyltransferase prephenate dehydrogenase | Amino Acid |
| PA3296 | *phoA* | Alkaline phosphatase | Cofactors and Vitamins |
| PA3333 | *fabH2* | 3-Oxoacyl-[acyl-carrier-protein] synthase III | Lipid |
| PA3633 | *ygbP* | 4-Diphosphocytidyl-2-C-methylerythritol synthase | Lipid |
| PA3659 | *dapC* | Succinyldiaminopimelate transaminase | Amino Acid |
| PA3686 | *adk* | Adenylate kinase | Nucleotide |
| PA4050 | *pgpA* | Phosphatidylglycerophosphatase A | Lipid |
| PA4693 | *pssA* | Phosphatidylserine synthase | Lipid |
| PA4770 | *lldP* | L-lactate permease | Transport |
| PA5322 | *algC* | Phosphomannomutase AlgC | Carbohydrate |
| PA5357 | *ubiC* | Chorismate pyruvate lyase | Cofactors and Vitamins |

In addition to changes in the GPR formulation for specific reactions, we also identified a potential change to the biomass reaction. Two PAO1 genes, *glgA* (PA2165) and *algC* (PA5322), are incorrectly predicted as essential for the synthesis of glycogen, a biomass component. Glycogen is not an essential metabolite for *P. aeruginosa* growth; however, it is very important for energy storage, which is why it was initially included in the biomass reaction [29]. Removal of glycogen from the biomass equation would make *glgA* and *algC* accurate predictions as nonessential genes in PAO1. Implementing these proposed changes in the PAO1 and PA14 GENREs resulted in enhanced predictive capability of the models (Dataset_S10, Dataset_S11, Table S3). The updated PAO1 model predicted consensus gene essentiality status in LB media with an accuracy of 97.4% compared to 96.8% for the original model. Meanwhile, the updated PA14 model predicted consensus gene essentiality status in LB media with an accuracy of 90.5% compared to 90.0% for the original mode. It is worth noting that, although these changes to the reconstructions were made to address essentiality discrepancies in LB media conditions, they also improved the PAO1 model predictive capabilities for consensus genes in sputum media, increasing accuracy from 92.6% to 93.0%.

While we identified several changes to the model to improve predictions, there were several genes for which we could find no literature evidence to change their predicted essentiality status. These genes highlight gaps in our current knowledge and understanding of *Pseudomonas* metabolism and indicate areas of future research.

Identification of these knowledge gaps is not possible without the reconciliation of experimental data with model predictions. Ultimately, this analysis demonstrates the utility of integrating data with GENREs to improve gene annotation and suggest areas of future research.

In addition to contextualizing essentiality for a given media condition, we also used the model to explain why certain metabolic genes are essential in one media-type versus another. We compared consensus LB essential genes to consensus sputum essential genes for PAO1 and identified the essential genes that were either shared by both conditions or unique to one condition versus the other. Overall, 18 genes were commonly essential, while 92 genes were uniquely essential in sputum and 26 genes were uniquely essential in LB, indicating the presence of condition-dependent essential genes.

We then focused our analysis just on those genes that were also present in the PAO1 model and compared these lists to model predictions. We found four genes that both the model and the screens indicated as uniquely essential in sputum but not in LB. Interestingly, all four of these genes (*pyrB, pyrC, pyrD*, and *pyrF*) are involved in pyrimidine metabolism. Applying flux sampling [30] to the PAO1 metabolic network model, we investigated why these four genes were uniquely essential in sputum but not in LB (Figure 2E). The pyrimidine metabolic pathway directly leads to the synthesis of several key biomass precursors (UMP, CMP, dCMP and dTMP), making it an essential subsystem within the network. Under LB media conditions, there are two inputs into the pathway, one through L-Glutamine and the other through Cytosine. However, in sputum media conditions, L-Glutamine is the only input into the pathway. Because of this reduction in the number of available substrates in sputum media, the steps for L-Glutamine breakdown must be active to synthesize the biomass precursors. Thus, the genes responsible for catalyzing this breakdown are essential in sputum media conditions. In contrast, because there are two LB substrates that feed into pyrimidine metabolism, if a gene involved in the breakdown of one of the substrates is not functional the other substrate is still accessible, thus making the deletion of that gene nonessential.

As stated above, further constraining the model with profiling data from both media conditions would help to further contextualize differences in the essentiality results by modulating the availability of certain reactions. Nevertheless, as demonstrated here, the metabolic network reconstruction can be a useful tool for providing functional explanations for why certain genes are essential in one condition versus another.

### Quantitative evaluation of the impact of media formulation on condition-independent essential gene identification

Given the variability in the number of candidate essential genes across the screens, we were interested in using the models to quantitatively evaluate the impact of media conditions on essentiality. We first focused our analysis on how the number of considered minimal media conditions impacts the number of condition-independent essential genes identified, or the number of genes found as essential in every condition. To do this, we simulated growth of the PA14 model on 42 different minimal media and performed *in silico* gene knockouts, identifying the genes essential for biomass production on each media condition (Figure 3A). We then randomly selected groups of minimal media conditions and compared their essential gene lists to determine the commonly essential genes, defined as the overlap. We performed this random selection of minimal media conditions for group sizes ranging from two to 40 minimal media conditions considered. For each group size, we randomly selected minimal media conditions 500 times. As expected, the more media conditions considered, the smaller the overlap of essential genes (Figure 3B). This relationship between the number of media conditions considered and the size of the overlap is best characterized by an exponential decay, with the size of the overlap eventually converging on 131 genes as 40 conditions are considered. This result suggests that to identify a core set of condition-independent essential genes, dozens of minimal media screens need to be compared. However, variability between the screens, as indicated by the error bars, could still confound interpretation, necessitating the comparison of replicates and potentially even more screens to truly identify condition-independent essential genes with high confidence.

**Figure 3.**
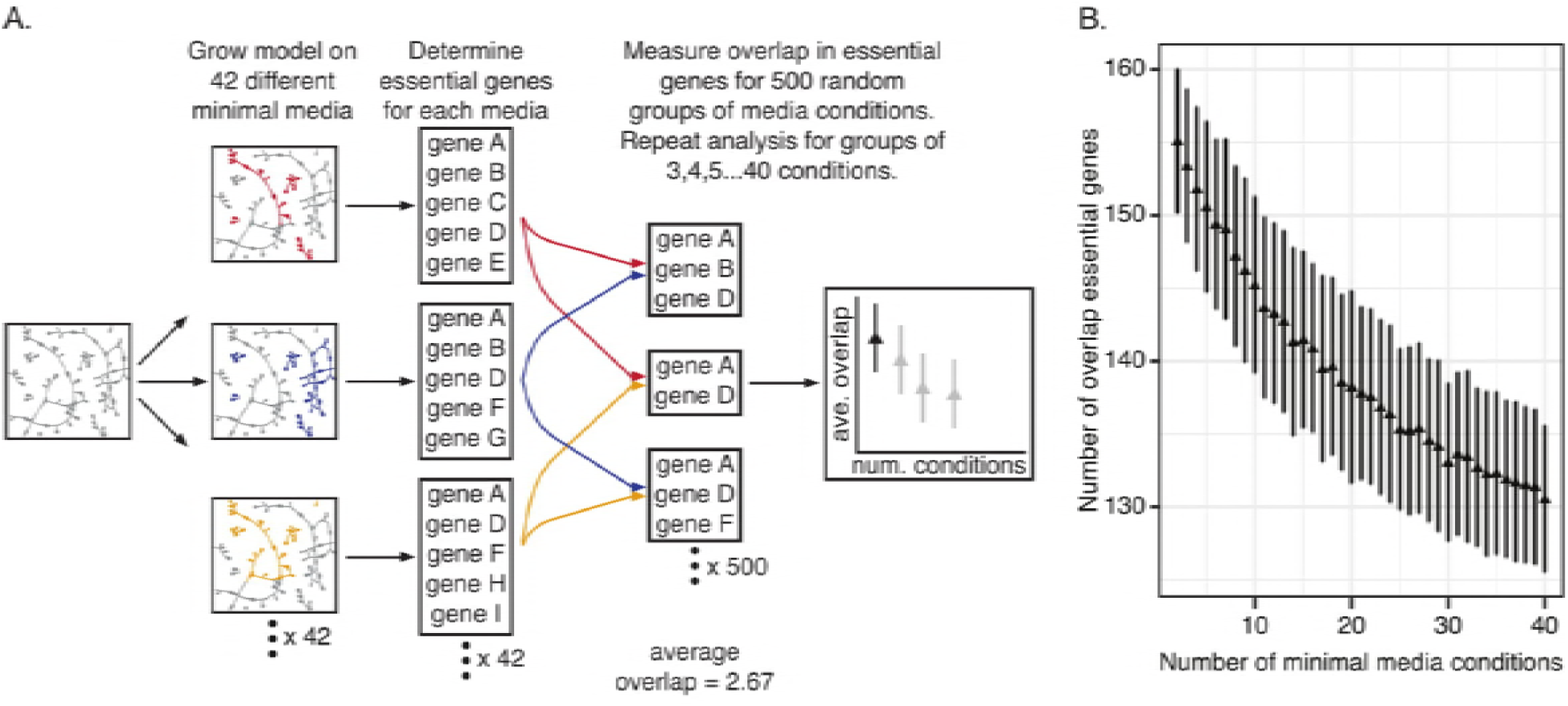
Computational assessment of the impact of number of minimal media conditions considered on condition-independent essentiality. (A) Pipeline for computational assessment of the impact of minimal media composition on condition-independent essentiality. The base PA14 model is grown on 42 different minimal media. For each minimal media condition, the *in silico* essential genes are identified, resulting in 42 essential gene lists. Initially, pairwise comparisons are made between minimal media essential gene lists to identify the shared essential genes. Specifically, the essential gene lists from two randomly selected minimal media conditions are compared to determine the overlap between the two gene lists. This random selection of two minimal media conditions to compare is repeated 500 times. The average number of overlap genes for all 500 comparisons is calculated as well as the standard deviation. Ultimately, this random selection of groups of minimal media conditions to compare is repeated for groups of three minimal media conditions, groups of four, and so on, up to groups of 40 minimal media conditions. (B) Impact of minimal media differences on the identification of condition-independent essential genes. Each data point represents the mean from 500 comparisons. Error bars indicate standard deviation.

We next assessed how modifications to a rich media, like LB, impact gene essentiality. LB is a complex media with known batch-to-batch variability [31,32], motivating this analysis of how differences in LB composition can alter essentiality. Given the challenge of modeling concentration, here the simulations focus on the presence or absence of metabolites in LB media. Specifically, we randomly selected carbon source components from LB media in sets of varying sizes, ranging from two to 21 LB media components considered. We then used these sets as the model media conditions and performed *in silico* gene knockouts to identify essential genes for biomass production on each LB media formulation (Figure 4A). For each set size, we randomly selected LB components 100 times and calculated the average number of essential genes identified as well as the number of shared essential genes across all 100 sets. As the number of LB media components increases, we found that the size of the essential gene lists decreases linearly (Figure 4B). If we were to consider even more media components beyond the scope of LB, we predict that this linear relationship would eventually plateau due to limitations in the metabolic network. This result suggests that a media richer than LB may be necessary to identify a core set of condition-independent essential genes.

**Figure 4.**
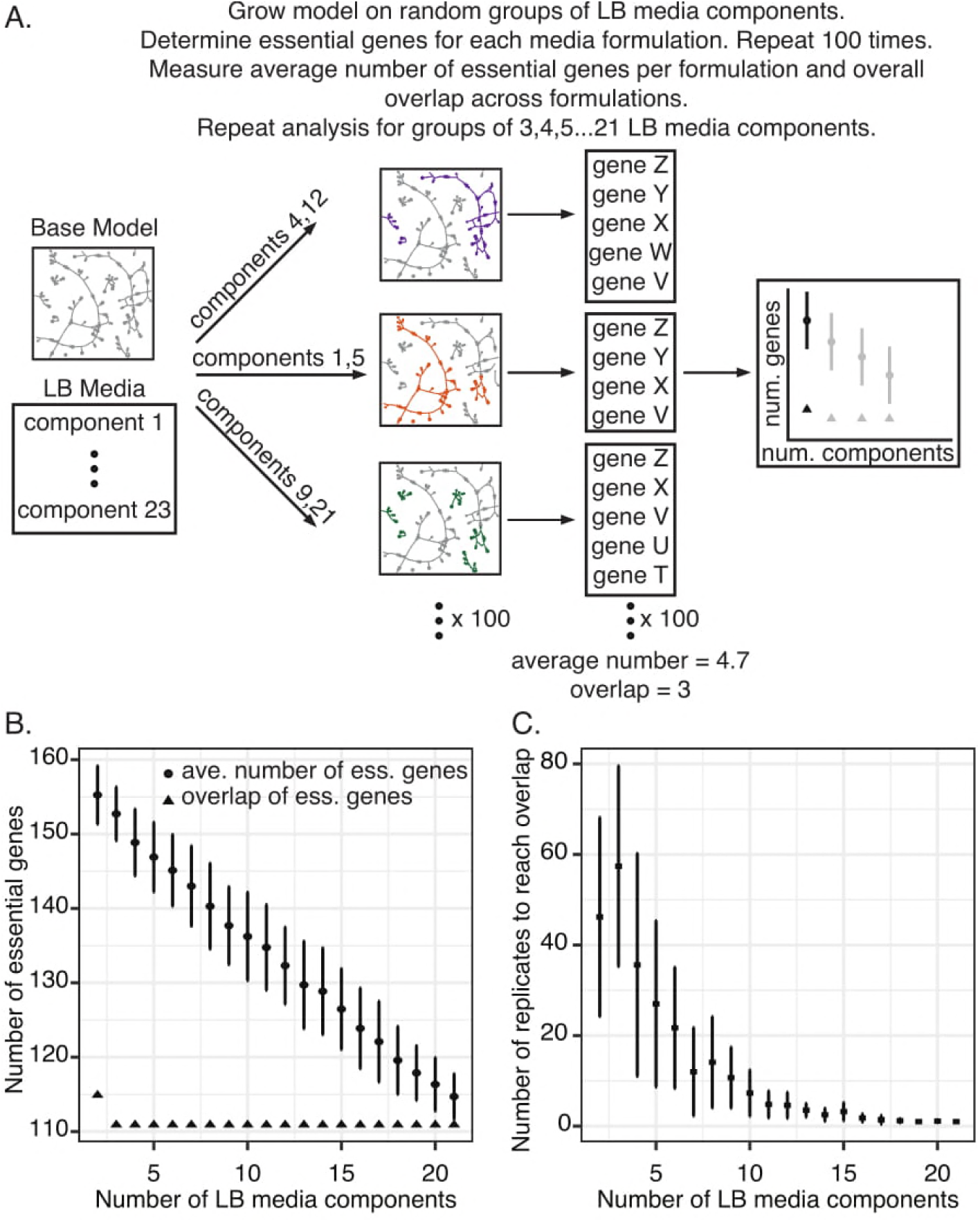
Computational assessment of the impact of LB media composition on condition-independent essentiality. (A). Pipeline for computational assessment of the impact of LB media formulation on condition-independent essentiality. The PA14 model is grown on different media formulations consisting of random groups of LB components. For instance, two random LB components are selected out of a pool of 23 LB components. The model is grown on these randomly selected pairs and the essential genes for growth on this media formulation are identified. This analysis is repeated 100 times for 100 pairs of LB media components. The average number of essential genes for growth on these random pairs across 100 different formulations is calculated as well as the standard deviation. Additionally, the essential genes common to all 100 different formulations is determined. Ultimately, this random selection of groups of LB media components to support growth of the model and essential gene identification is repeated for groups of three LB components, groups of four, and so on, to groups of 21 LB media components. (B) Impact of LB media formulation on the identification of condition-independent essential genes. Circles represent the average number of essential genes identified in different LB media formulations across 100 comparisons. Triangles represent the shared essential genes (i.e., the overlap) across all 100 comparisons. Error bars indicate standard deviation. (C) Number of replicates needed to converge on shared essential genes in different LB formulations. The pipeline outlined in Panel A was repeated 10 independent times, with 100 replicates per set size. For each iteration, the number of replicates needed to recapture the 111 overlapping genes was calculated. Each data point represents the average number of replicates from the 10 runs. Error bars indicate standard deviation.

Interestingly, we found that as more complex LB media formulations are considered, the number of shared essential genes across 100 simulations quickly converges on 111. Indeed, only three LB media components were needed to achieve this overlap. Thus, even though the average size of essential gene lists is larger for less complex media formulations, the overlap of these larger essential gene lists still results in the same overlap as more complex media formulations, suggesting that changes in complex media formulation have minimal impact on determining a core set of essential genes.

However, for this analysis, we had compared 100 random media formulations for each set size, potentially masking the impact of media changes on essentiality. To identify how many LB media formulations need to be compared to converge on this overlap value, we re-ran this analysis 10 times and, for each iteration, determined the number of samples, or replicates, needed to recapture the 111 overlapping genes (Figure 4C). In more complex media formulations, relatively few comparisons are needed to identify the 111 overlapping essential genes. However, as fewer LB media components are considered, more comparisons need to be made. For example, in the case of formulations consisting of only three LB media components, nearly 60 comparisons are needed to converge on the 111 overlap essential genes. Thus, as the media formulation diverges from true LB due to batch-to-batch variability, more comparisons are necessary to converge on a core set of essential genes.

Taken together, these computational analyses define the scope that is needed to identify condition-independent essential genes. These results suggest that both the number of media conditions and the number of replicates analyzed can impact our ability to determine condition-independent essential genes.

## Discussion

The identification of both condition-dependent and condition-independent essential genes has been a long-standing interest [33,34]. Determination of these essential processes can aid in the discovery of novel antibacterial targets as well as the discovery of minimal genomes required to sustain life [7,35]. In this study, we performed a large-scale comparison of multiple gene essentiality datasets and contextualized essential genes using genome-scale metabolic network reconstructions. We applied this approach to several *P. aeruginosa* transposon mutagenesis screens performed on multiple media conditions and demonstrated the utility of GENREs in providing functional explanations for essentiality and resolving differences between screens. Finally, using the *P. aeruginosa* GENRE, we performed a high-throughput, quantitative analysis to determine how media conditions impact the identification of condition-independent essential genes. The resulting insights would be challenging to develop without the use of a computational model of *P. aeruginosa* metabolism. Our work enables the elucidation of mechanistic explanations for essentiality, which is challenging to determine experimentally. Ultimately, this approach serves as a framework for future contextualization of gene essentiality data and can be applied to any cell type for which such data is available. Additionally, by quantifying the impact of media conditions on the identification of condition-independent essential genes, we contribute novel insights for design of future gene essentiality screens and identification of core metabolic processes.

Recent advances in deep-sequencing technologies combined with transposon mutagenesis have enabled high-throughput determination of candidate essential genes for a variety of bacterial species in a wide range of environmental conditions [36]. While researchers have demonstrated reasonable reproducibility within a given study [37], variability across studies has been suggested but not assessed on a large-scale [1,38]. Our comparison of multiple *P. aeruginosa* transposon mutagenesis screens revealed substantial variability in candidate essential genes within and across media conditions, particularly for strain PAO1. Numerous factors may contribute to this lack of overlap between the screens, such as differences in transposon insertion library complexity, differences in data analysis and statistical determination of essentiality, as well as environmental variability between the screens [8,9]. Factors such as these lead to discrepancies between screens and complicate our ability to identify high-confidence sets of condition-dependent and condition-independent essential genes.

Focusing on one of these factors, we used the metabolic model of *P. aeruginosa* strain PA14 to quantitatively assess how media formulation impacts the identification of condition-independent essential genes. While previous *in vitro* studies have surveyed conditional essentiality in numerous environmental conditions, these screens used an already established mutant library for each media-type [39]. In this work, we computationally generated *de novo* mutant libraries for individual media conditions, eliminating any bias from starting with an established mutant library. Ultimately, we found that to determine a high-confidence set of core essential genes for minimal media conditions, more than 40 minimal media formulations need to be compared. We extended this analysis to consider how differences in rich media formulations impact gene essentiality and found that as rich media formulations diverge, as many as 60 replicates are needed to identify condition-independent essential genes with high-confidence. Taken together, these computational results suggest a rich opportunity for a large-scale experimental effort to identify with high confidence condition-independent essential genes. These insights would be impossible to garner without computational modeling due to the sheer number of comparisons made.

In addition to variability between datasets, a central difficulty of performing gene essentiality screens lies in the interpretation of why a gene is essential in a given condition. Oftentimes, laborious follow-up experiments are necessary to investigate the role of a gene in a given condition using lower-throughput approaches [36]. Here, we presented a strategy for contextualizing gene essentiality data using genome-scale metabolic network reconstructions. We demonstrated the utility of this approach by providing functional reasons for essentiality for consensus LB media essential genes. For these genes, we determined which specific components of biomass could not be synthesized when the gene was knocked out. Additionally, by analyzing the network structure and flux patterns, we used the model to explain why certain genes are essential in one condition versus another. Our computational approach provides testable hypotheses regarding the functional role of a gene in synthesizing biomass in a given environmental condition, streamlining downstream follow-up experiments. In future work, profiling data could be integrated with the metabolic networks to further enhance the utility of these models in contextualizing gene essentiality [24]. Additionally, integration of transcriptional regulatory networks with the GENREs would further expand the number of genes considered [40].

In summary, genome-scale metabolic network reconstructions can guide the design of gene essentiality screens and help to interpret their results. The identification of both condition-independent and condition-dependent essential genes is vital for the discovery of novel therapeutic strategies and mechanistic modeling streamlines the ability to identify these genes. This framework can be applied to numerous other organisms of both clinical and industrial relevance.

## Acknowledgments

The authors would like to thank Jennifer Bartell and Joanna Goldberg for early discussions related to the project, Laura Dunphy, Kevin Janes, and Phillip Yen for thoughtful comments on the manuscript, and Sean Leonard for assistance in analysis of the transposon sequencing raw datasets. Support for this project was provided by The Wagner Fellowship and Unilever.

## Author Contributions

A.S.B. and J.A.P. conceived and designed the study. A.S.B. completed all analyses. A.S.B. and J.A.P. wrote and edited the manuscript.

## Methods

### Data sources

Transposon insertion library datasets were downloaded from the original publication for each screen where available. Screens were renamed following this pattern: *Strain.Media.NumEssentials*, where *Strain* indicated whether the screen was for strain PAO1 or PA14, *Media* indicated which media condition the screen was performed on, and *NumEssentials* indicated the number of essential genes identified for the given strain on the given media condition. Specifically, for the PAO1.LB.201, PAO1.Sputum.224, and PAO1.Pyruvate.179 datasets, Dataset_S01 was downloaded from [19]. For the PAO1.LB.335, PAO1.Sputum.405, and PAO1.Succinate.640 datasets, Dataset_S01 was downloaded from [18]. For the PA14.LB.634 dataset, Table S1 was downloaded from [17]. For the PA14.Sputum.510 dataset, Dataset_S04 was downloaded from [18]. For the PAO1.LB.913 dataset, PA_two_allele_library5.xlsx was downloaded from the Manoil Laboratory website (http://www.gs.washington.edu/labs/manoil/libraryindex.htm). For the PA14.LB.1544 dataset, NRSetFile_v5_061004.xls was downloaded from the PA14 Transposon Insertion Mutant Library website (http://pa14.mgh.harvard.edu/cgibin/pa14/downloads.cgi).

The PAO1 and PA14 genome-scale metabolic network reconstructions were downloaded from the Papin Laboratory website (http://www.bme.virginia.edu/csbl/Downloads1-pseudomonas.html).

### Generation of candidate essential gene lists

Candidate essential genes were determined for each screen as follows. For PAO1.LB.201, we considered genes to be essential if they were not disrupted in all six of the Tn-seq runs on LB in the original dataset. For PAO1.Sputum.224, we considered genes to be essential if they were not disrupted in all four of the Tn-seq runs on sputum in the original dataset. For PAO1.Pyruvate.179, we considered genes to be essential if they were not disrupted in all three of the Tn-seq screens on Pyruvate minimal media in the original dataset. For PAO1.LB.335, PAO1.Sputum.405, and PAO1.Succinate.640, we used the genes that were labeled as essential in the original dataset. For PAO1.LB.913, the mutants listed in the transposon insertion library were compared to a list of all known genes in the PAO1 genome. Genes in the PAO1 genome that were not in the mutant library list were considered to be essential. For PA14.LB.634, we used the genes listed as essential in the original dataset. For PA14.BHI.424 and PA14.Sputum.510, we used the genes that were labeled as essential in the original dataset. For PA14.LB.1544, the mutants listed in the transposon insertion library were compared to a list of all known genes in the PA14 genome. Genes in the PA14 genome that were not in the mutant library list were considered to be essential.

### Comparison of candidate essential gene lists

Hierarchical clustering with complete linkage was performed on the candidate essential gene lists for the PA14 and PAO1 screens and visualized with a dendrogram. The overlap between the datasets was visualized using the R-package, UpsetR [41].

### Re-analysis of transposon sequencing datasets

PAO1.LB.335 sequencing data were downloaded from NCBI SRA under the accession number SRX031647. PAO1.LB.201 sequencing data were downloaded from NCBI SRA under the accession number PRJNA273663. Data were analyzed using methods adapted from [18,20]. Briefly, reads were mapped to the PAO1 reference genome (GCA_000006765.1 ASM676v1 assembly downloaded from NCBI) using bowtie2 v.2.3.4.1. Open reading frame assignments were modified where 10% of the 3’ end of every gene was removed in order to disregard insertions that may not interrupt gene function. Aligned reads were mapped to genes and we removed the 50 most abundant sites to account for potential PCR amplification bias. We applied weighted LOESS smoothing to correct for genome position-dependent effects. One-hundred random datasets were generated by randomizing insertion locations. Previous analysis showed that results begin to converge after 50 random datasets [18]. We compared the random datasets to the experimental datasets with a negative binomial test in DESeq2. We corrected for multiple testing by adjusting the p-value with the Benjamini-Hochberg method. We used the mclust package in R to test whether a gene was ‘reduced’ or ‘unchanged’. Genes were called ‘essential’ if they were assigned to the ‘reduced’ category by mclust with an adjusted p-value <0.05 and uncertainty <0.1.

### Model gene essentiality predictions

*In silico* gene essentiality screens were performed in relevant media conditions using the PAO1 and PA14 genome-scale metabolic network reconstructions [23]. Specifically, media formulations were computationally approximated for LB, sputum, pyruvate minimal media, and succinate minimal media for the PAO1 simulations and LB and sputum for the PA14 simulations. Systematically, genes were deleted from the models one-by-one and the resulting impact on biomass production was assessed. If biomass production for the associated mutant model was below 0.0001 h^-1^, a standard threshold, the knocked-out gene was predicted to be essential [23]. For each *in silico* predicted essential gene, we determined which biomass components specifically could not be synthesized using the COBRA toolbox function, biomassPrecursorCheck() [42]. Statistical significance for the comparison of the “mismatch: model nonessential, screen essential” category and the “mismatch: model essential, screen nonessential” category was assessed using the Wilcoxon signed-rank test.

### Subsystem assignment of consensus essential and nonessential genes

For each of the consensus essential and nonessential genes that were also present in the PAO1 and PA14 models, we determined which subsystems they participated in using an in-house script (see Supplementary Information). Briefly, we first converted model subsystems to broad subsystems based on KEGG functional categories [43]. We then identified the reactions associated with the gene of interest and used the broad subsystem of this reaction to indicate the subsystem assignment for the gene of interest. Where there was more than one reaction connected to a gene, we used the reaction associated with the first instance of the gene in the network for subsystem assignment.

### Flux sampling in LB and sputum

The impact of media conditions on flux through pyrimidine metabolism in the PAO1 metabolic network reconstruction was assessed using the flux sampling algorithm optGpSampler [30]. Briefly, optGpSampler samples the solution space of genome-scale metabolic networks using the Artificial Centering Hit-and-Run algorithm and returns a distribution of possible flux values for reactions of interest. Three-thousand flux samples were collected for each simulation, using one thread and a step-size of one. Maximization of biomass synthesis was set as the objective function. Flux sampling simulations were performed for PAO1 grown in LB media and sputum media. The median flux values for every reaction in pyrimidine metabolism were compared between the LB and sputum simulations to determine whether flux was higher, lower, or unchanged in sputum versus LB.

### Media formulation impact on essentiality

The impact of media formulation on gene essentiality predictions was assessed using the PA14 genome-scale metabolic network reconstruction. For the minimal media analysis, the PA14 model was grown on 42 different minimal media and *in silico* essential genes were identified as described above. We then randomly selected groups of minimal media conditions of varying sizes, ranging from two to 41 minimal media conditions considered, and found the intersection of the group’s predicted essential gene lists, or the genes that were identified as essential in every condition considered within that group. For each group size, we randomly selected minimal media conditions 500 times.

For the LB media analysis, we randomly selected components from LB media in sets of varying sizes, ranging from two to 21 LB media components considered, used these sets as the model media conditions, and identified *in silico* essential genes as above. For each set size, we randomly selected LB components 100 times and calculated the average total number of essential genes identified and the intersection of the essential genes across all 100 sets. To determine how many LB media formulations needed to be compared to converge on this intersection, we re-ran this LB media formulation analysis 10 times and, for each iteration, determined the number of samples needed to achieve the size of the overlap if all 100 samples were considered at each set size

### Code and data availability

Code and files necessary to recreate figures and data can be found here: https://github.com/ablazier/gene-essentiality

### Computational resources

The COBRA Toolbox 2.0.5 [42], the Gurobi 6.5 solver, and MATLAB R2016a were used for model simulations. optGPSampler1.1 was used for flux sampling simulations [30]. Bowtie2 v.2.3.4.1 [44] and Samtools v.1.3.1 [45] were used for transposon sequencing analysis. R 3.3.3 was used for all other analyses and figure generation.

## Supplementary Information

Dataset_S1.xls - PAO1 candidate essential genes for *in vitro* screens

Candidate essential genes lists for each PAO1 transposon mutagenesis screen. Candidate essential genes are marked with a ‘1’, while non-essential genes are marked with a ‘0’.

Dataset_S2.xls - PA14 candidate essential genes for *in vitro* screens

Candidate essential genes lists for each PA14 transposon mutagenesis screen. Candidate essential genes are marked with a ‘1’, while non-essential genes are marked with a ‘0’.

Dataset_S3.xls - PAO1 model predicted essential genes for *in silico* screens

Model predicted essential genes lists for PAO1 growth simulated on LB media, Sputum media, Pyruvate minimal media, and Succinate minimal media. Model predicted essential genes are marked with a ‘1’, while non-essential genes are marked with a ‘0’.

Dataset_S4.xls - PA14 model predicted essential genes for *in silico* screens

Model predicted essential genes lists for PA14 growth simulated on LB media and Sputum media. Model predicted essential genes are marked with a ‘1’, while non-essential genes are marked with a ‘0’.

Dataset_S5.xls - PAO1 consensus metabolic essential/non-essential genes

Lists of consensus metabolic essential and non-essential genes for PAO1 on LB media and Sputum media.

Dataset_S6.xls - PA14 consensus metabolic essential/non-essential genes

Lists of consensus metabolic essential and non-essential genes for PA14 on LB media.

Dataset_S7.xls - Biomass precursors for PAO1 model predicted consensus essential genes List of biomass precursors that cannot be synthesized when PAO1 model predicted consensus essential genes are removed from the model.

Dataset_S8.xls - Biomass precursors for PA14 model predicted consensus essential genes List of biomass precursors that cannot be synthesized when PA14 model predicted consensus essential genes are removed from the model.

Dataset_S9.xls - Proposed model changes

Table of proposed model changes based on discrepancies between model predictions and consensus metabolic non-essential genes for PAO1 on LB.

Dataset_S10.xls - PAO1 model predicted essential genes for *in silico* screens for the updated PAO1 model

Model predicted essential genes lists for PAO1 growth simulated on LB media and Sputum media. Model predicted essential genes are marked with a ‘1’, while non-essential genes are marked with a ‘0’.

Dataset_S11.xls - PA14 model predicted essential genes for *in silico* screens for the updated PA14 model

Model predicted essential genes lists for PA14 growth simulated on LB media. Model predicted essential genes are marked with a ‘1’, while non-essential genes are marked with a ‘0’.

Figure S1. Comparison of candidate essential genes from PAO1 LB transposon mutagenesis screens reveals variability across screens.

Figure S2. Comparison of candidate essential genes from LB transposon mutagenesis screens reveals variability across screens.

Figure S3. Distribution of PAO1 consensus essential and nonessential genes across model subsystems

Table S1. Detailed description of *in vitro* transposon mutagenesis screens.

Table S2. Percent accuracy between model predictions of essentiality and *in vitro* identified essential genes.

Table S3. Consensus metabolic essential and non-essential genes for PAO1 and PA14 media conditions with more than two screens.

---

**Figure S1.**
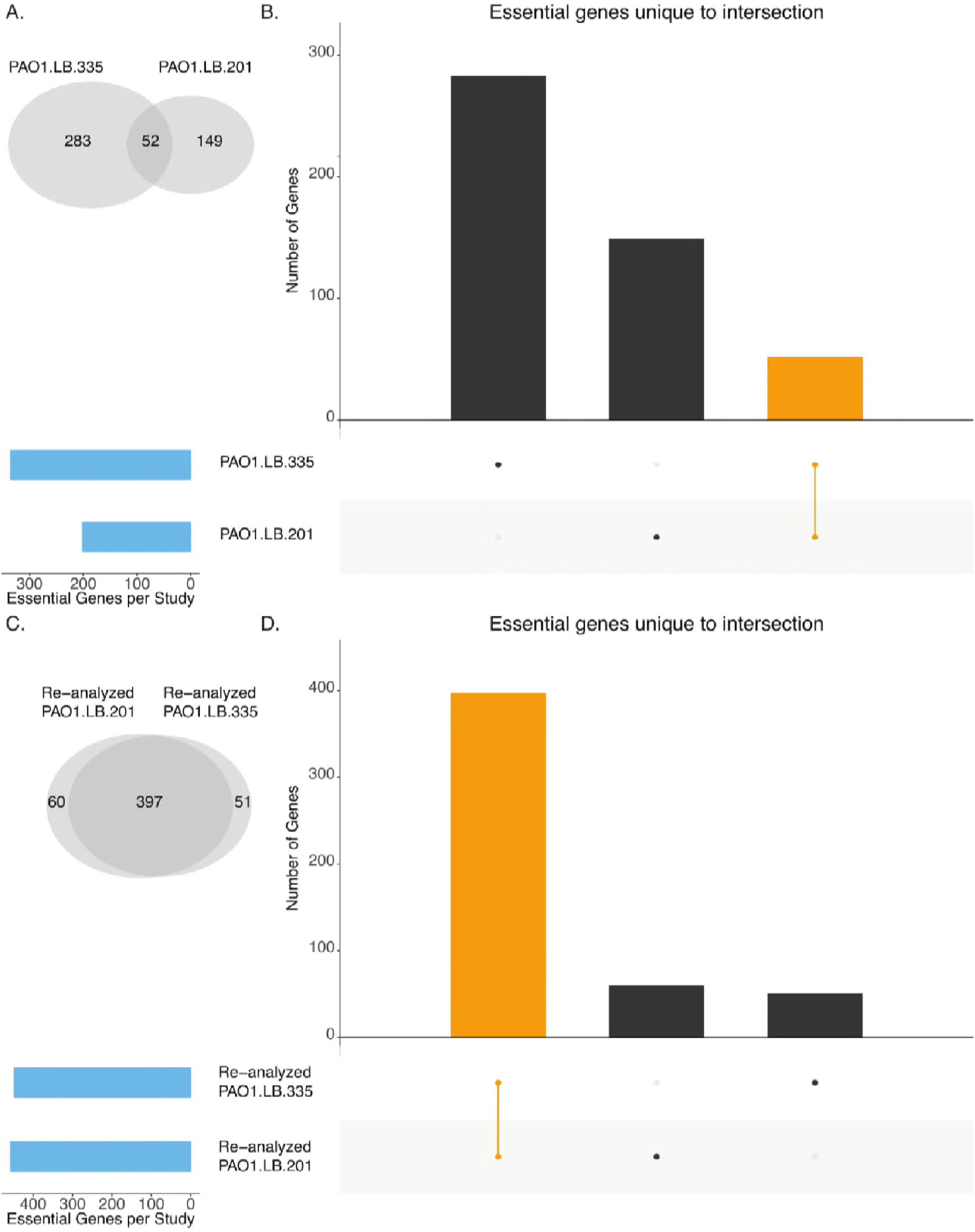
Comparison of candidate essential genes from PAO1 LB transposon mutagenesis screens reveals variability across screens. (A and C). Venn diagrams of original (Panel A) and re-analyzed (Panel C) candidate essential gene lists from PAO1 transposon mutagenesis screens performed on LB. (B and D). Overlap analysis of original (Panel B) and re-analyzed (Panel D) candidate essential gene lists for PAO1 transposon mutagenesis screens performed on LB. Blue bars indicate the total number of candidate essential genes identified in each screen. Black bars indicate the number of candidate essential genes unique to the intersection given by the filled-in dots. The orange bar indicates the overlap of both screens.

**Figure S2.**
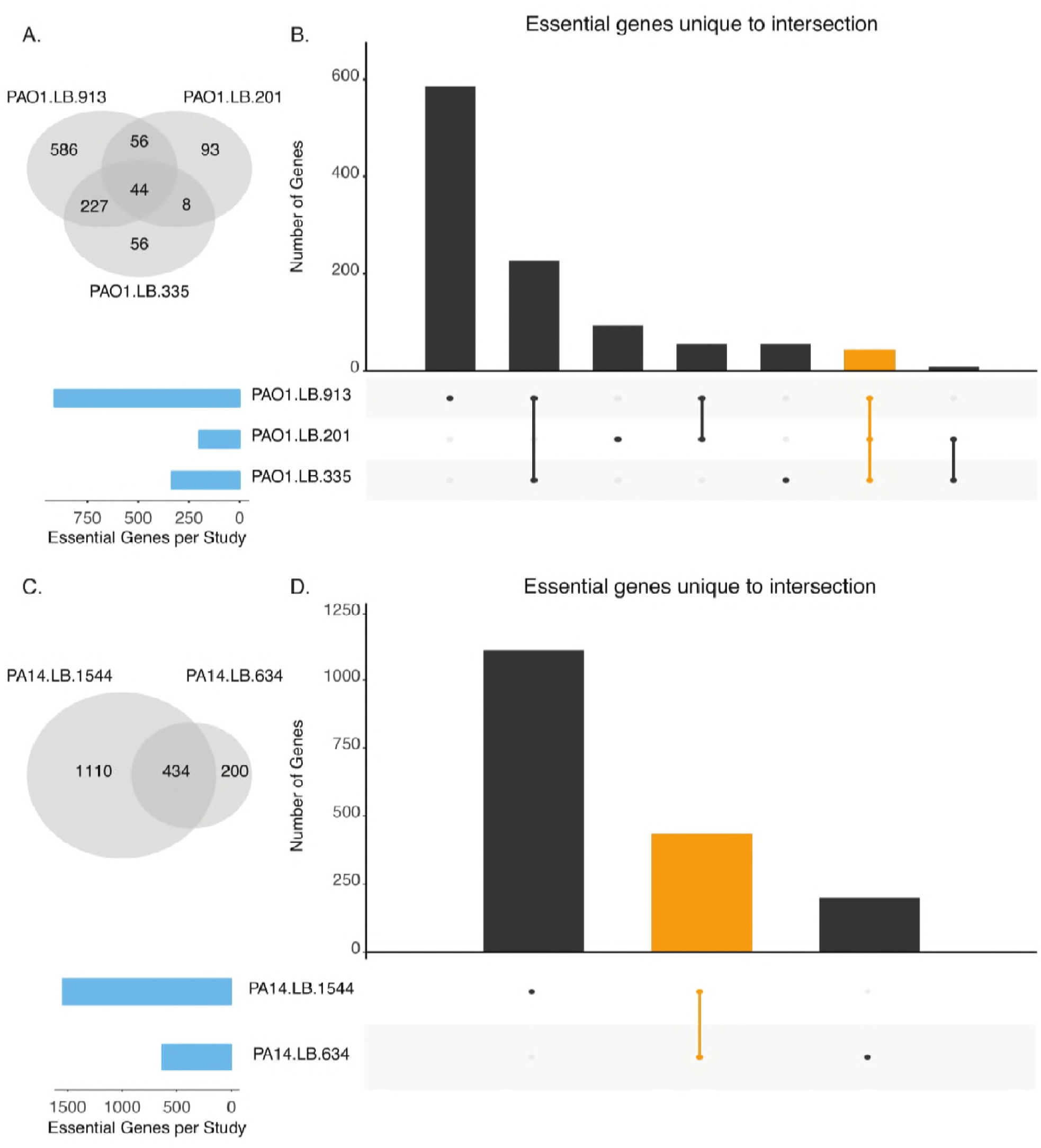
Comparison of candidate essential genes from LB transposon mutagenesis screens reveals variability across screens. (A and C). Venn diagram of candidate essential genes lists for transposon mutagenesis screens performed on LB for PAO1 and PA14, respectively. (B and D). Overlap analysis of candidate essential gene lists for transposon mutagenesis screens performed on LB for PAO1 and PA14, respectively. Blue bars indicate the total number of candidate essential genes identified in each screen. Black bars indicate the number of candidate essential genes unique to the intersection given by the filled-in dots. The orange bar indicates the overlap for all screens for either PAO1 (Panel B) or PA14 (Panel D). The black and orange bars correspond to the intersections identified in the venn diagrams in panels A and C.

**Figure S3.**
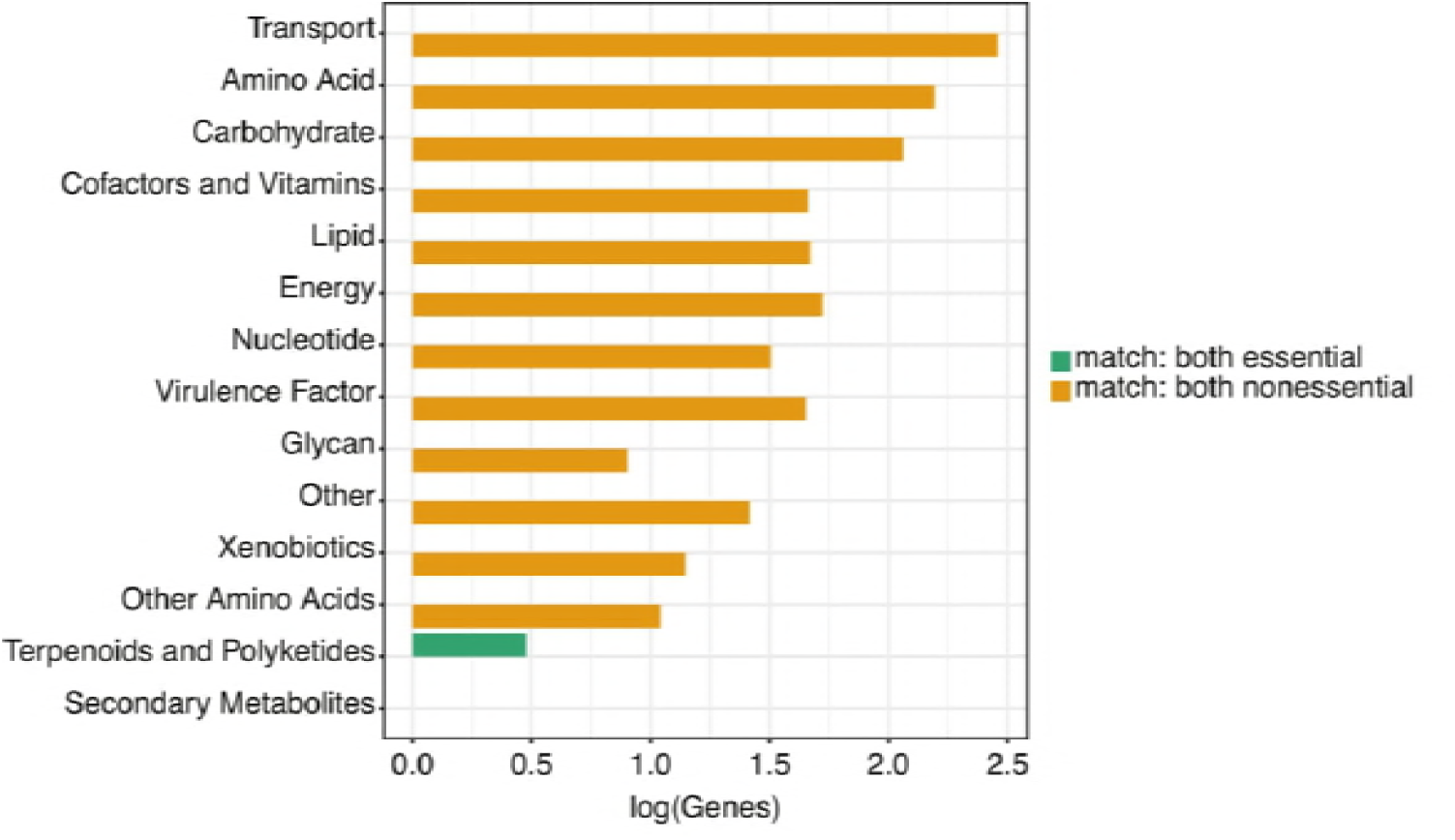
Distribution of PAO1 consensus essential and nonessential genes across model subsystems. Functional subsystems for PAO1 consensus essential and nonessential genes that were also identified to be essential or nonessential in the PAO1 genome-scale metabolic network model. Consensus essential and nonessential genes were identified for PAO1 by comparing all three LB screens and identifying those genes which were either essential or nonessential in all three screens.

**Table S1.** Detailed description of *in vitro* transposon mutagenesis screens.

| Screen | Strain | Media | Number of Essential Genes | Publication of analyzed datasets | Initial publication of dataset | Publication of initial insertion library | Experimental overview to generate initial insertion library | Analysis overview |
| --- | --- | --- | --- | --- | --- | --- | --- | --- |
| PAO1.LB.913 | PAO1 | LB | 913 | Jacobs et al., 2003 | Jacobs et al., 2003 | Jacobs et al., 2003 | Random transposon insertion mutagenesis with ISphoA and ISlacZ | Manual tally of recovered genes |
| PAO1.LB.201 | PAO1 | LB | 201 | Lee et al., 2015 | Lee et al., 2015 | Lee et al., 2015 | Tn-seq circle method with Tn5-based transposon T8 | Number of transposon insertions per gene compared to a normal distribution |
| PAO1.LB.335 | PAO1 | LB | 335 | Turner et al., 2015 | Gallagher et al., 2011 | Gallagher et al., 2011 | Tn-seq circle method with Tn5-based transposon T8 | Number of transposon insertions per gene compared to Monte Carlo simulated data |
| PAO1.Sputum.224 | PAO1 | Sputum | 224 | Lee et al., 2015 | Lee et al., 2015 | Lee et al., 2015 | Tn-seq circle method with Tn5-based transposon T8 | Number of transposon insertions per gene compared to a normal distribution |
| PAO1.Sputum.405 | PAO1 | Sputum | 405 | Turner et al., 2015 | Turner et al., 2015 | Gallagher et al., 2011 | Tn-seq circle method with Tn5-based transposon T8 | Number of transposon insertions per gene compared to Monte Carlo simulated data |
| PAO1.Pyruvate.179 | PAO1 | Pyruvate minimal media | 179 | Lee et al., 2015 | Lee et al., 2015 | Lee et al., 2015 | Tn-seq circle method with Tn5-based transposon T8 | Number of transposon insertions per gene compared to a normal distribution |
| PAO1.Succinate.640 | PAO1 | Succinate minimal media | 640 | Turner et al., 2015 | Turner et al., 2014 | Gallagher et al., 2011 | PCR-based Tn-seq method with Tn5-based transposon T8 | Number of transposon insertions per gene compared to Monte Carlo simulated data |
| PA14.LB.1544 | PA14 | LB | 1544 | Liberati et al., 2006 | Liberati et al., 2006 | Liberati et al., 2006 | Random transposon insertion mutagenesis with mariner family transposon | Manual tally of recovered genes |
| PA14.LB.634 | PA14 | LB | 634 | Skurnik et al., 2013 | Skurnik et al., 2013 | Skurnik et al., 2013 | INSeq method with mariner family transposon | Fold change between reads per kilobase per million reads |
| PA14.Sputum.510 | PA14 | Sputum | 510 | Turner et al., 2015 | Turner et al., 2015 | Turner et al., 2015 | PCR-based Tn-seq method with Tn5-based transposon T8 | Number of transposon insertions per gene compared to Monte Carlo simulated data |

**Table S2.** Percent accuracy between model predictions of essentiality and *in vitro* identified essential genes.

| Screen | % Accuracy |
| --- | --- |
| PAO1.LB.913 | 87.46 |
| PAO1.LB.201 | 84.76 |
| PAO1.LB.335 | 89.29 |
| PAO1.Sputum.224 | 84.32 |
| PAO1.Sputum.405 | 87.63 |
| PAO1.Pyruvate.179 | 86.07 |
| PAO1.Succinate.640 | 83.28 |
| PA14.LB.1544 | 79.95 |
| PA14.LB.634 | 84.63 |
| PA14.Sputum.510 | 87.81 |

**Table S3.**
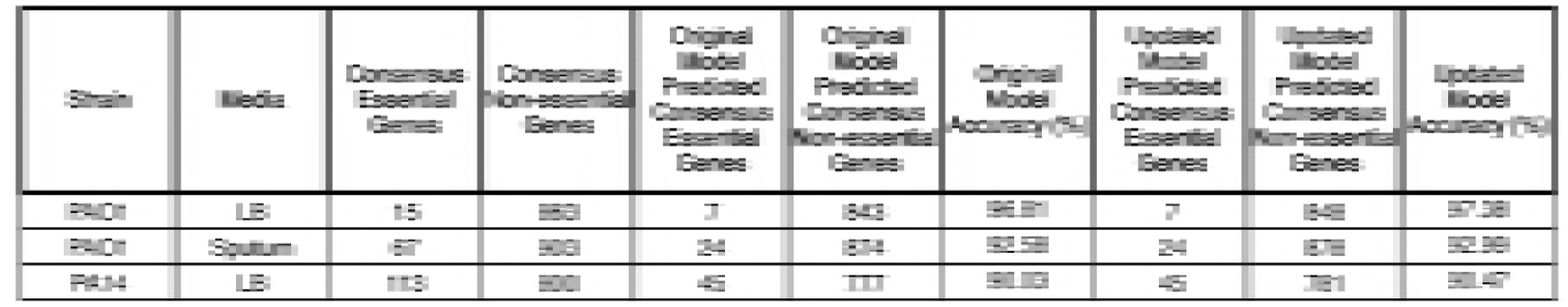
Consensus metabolic essential and non-essential genes for PAO1 and PA14 media conditions with more than two screens.

